# Evolution of herbicide resistance through co-option of an ethylene-associated nitrilase

**DOI:** 10.64898/2026.11.06.764846

**Authors:** Hinata Ishizawa, Luna Nakanomori, Kowit Hengphasatporn, Kazusa Nishimura, Masumi Ishizaka, Chiharu Akimoto-Tomiyama, Danrui Su, Manami Sugahara, Masaki Endo, Rintaro Suzuki, Mitsuhiko P. Sato, Kentaro Yasuda, Yasuteru Shigeta, Takahiro Gondo, Akira Uchino, Satoshi Iwakami

**Affiliations:** Graduate School of Agriculture, Kyoto University, Kitashirakawa-Oiwake-cho, Kyoto 606-8502 JAPAN; Graduate school of Agriculture, Tokyo University of Agriculture and Technology, Fuchu, Tokyo 183-8509 JAPAN; Center for Computational Sciences, University of Tsukuba, Tsukuba 305-8573 JAPAN; Graduate School of Environmental, Life, Natural Science and Technology, Okayama University, Okayama 700-8530 JAPAN; Research Center for Advanced Analysis, National Agriculture and Food Research Organization, Tsukuba, Ibaraki 305-8518 JAPAN; Institute of Agrobiological Sciences, National Agriculture and Food Research Organization, Tsukuba, Ibaraki 305-8634 JAPAN; United Graduate School of Agricultural Science, Tokyo University of Agriculture and Technology, Fuchu, Tokyo 183-8509 JAPAN; Faculty of Agriculture, University of Miyazaki, Miyazaki 889-2192 JAPAN; Kazusa DNA Research Institute, Chiba 292-0818 JAPAN; Agri-Innovation Education and Research Center, Akita Prefectural University, Akita 010-0451 JAPAN; Frontier Science Research Center, University of Miyazaki, Miyazaki 889-2192 JAPAN; Central Region Agricultural Research Center, National Agriculture and Food Research Organization, Tsu, Mie 514-2392 JAPAN; Institute of Agriculture, Tokyo University of Agriculture and Technology, Fuchu, Tokyo 183-8509 JAPAN

## Abstract

Adaptive evolution frequently proceeds through the co-option of pre-existing biological functions, yet the molecular mechanisms by which such co-option generates new adaptive traits over contemporary evolutionary timescales remain poorly understood. Herbicide-resistant weeds provide a powerful model for addressing this question because enhanced herbicide metabolism has repeatedly evolved within only a few decades. Here, we show that resistance to cyhalofop-butyl in *Echinochloa crus-galli* evolved through the co-option of *EcNIT4A2*, which encodes a nitrilase that normally functions in ethylene-associated cyanide detoxification. Genetic mapping, transcriptomics and transgenic analyses identified *EcNIT4A2* as the major determinant of resistance. A D317G substitution enhanced its herbicide-detoxifying activity. Molecular dynamics simulations indicated that this substitution facilitates substrate access to the catalytic pocket. A promoter structural variant markedly increased *EcNIT4A2* expression. Our findings reveal how coding and regulatory changes can jointly drive the co-option and optimization of a conserved metabolic function during rapid adaptation under intense anthropogenic selection.

## Introduction

Adaptive evolution frequently proceeds through the modification and redeployment of pre-existing biological systems rather than the evolution of entirely new ones.^1,2^ In plants, evolutionary innovation is repeatedly linked to gene duplication and the subsequent co-option of existing genes into new functional contexts.^3^ Such co-option may be particularly crucial for traits that evolve over contemporary timescales, where adaptation must arise within very few generations. However, the molecular processes driving evolutionary co-option over such short timescales remain largely unknown.

Herbicide-resistant weeds pose an escalating threat to sustainable agriculture and global food security by compromising the efficacy of essential weed control tools.^4–6^ Beyond its agricultural importance, the repeated and independent evolution of herbicide resistance under exceptionally strong anthropogenic selection provides an outstanding model for investigating contemporary adaptive evolution.^7,8^ One common outcome of this selection is the evolution of metabolic herbicide resistance through enhanced detoxification capacity.^4,9^

Unlike mammals, in which several enzyme families have specialized roles in xenobiotic metabolism, plants do not appear to possess enzyme families dedicated primarily to detoxification of foreign chemicals.^10^ Thus, herbicide-metabolizing enzymes are generally believed to have evolved from enzymes whose primary functions lie in endogenous or specialized metabolism.^11–15^ Nevertheless, research into metabolic resistance has largely focused on a limited set of enzyme families already known for their capacity to metabolize xenobiotics, particularly cytochrome P450 monooxygenases and glutathione S-transferases.^16,17^ Whether enzymes with established roles in endogenous metabolism can also be recruited for herbicide detoxification therefore remains largely unexplored.

Here we investigate the molecular basis of resistance to the acetyl-CoA carboxylase (ACCase) inhibitor cyhalofop-butyl (CB) in the globally problematic grass weed *Echinochloa crus-galli*.^18^ We identify EcNIT4A2, a NIT4 nitrilase involved in ethylene-associated cyanide metabolism, as the major determinant of CB resistance. By integrating quantitative genetics, comparative transcriptomics, functional analyses and molecular dynamics simulations, we show that resistance arose through the co-option of this endogenous metabolic enzyme. A promoter structural variant (SV) markedly increased *EcNIT4A2* expression, whereas a single amino acid substitution enhanced its herbicide-detoxifying activity by facilitating substrate access to the catalytic pocket. Together, our findings reveal how regulatory and coding changes can jointly recruit and optimize a conserved metabolic function during rapid adaptation under intense anthropogenic selection.

## Results

### The R1 line rapidly detoxifies cyhalofop-acid to cyhalofop-diacid

Previous studies showed that two CB-resistant lines collected in 2010 from nearby rice fields in Japan exhibited distinct responses to different herbicides despite their close geographic proximity (Fig. 1a,b).^18,19^ Because neither line carried known target-site resistance mutations,^18^ we investigated whether enhanced herbicide metabolism underlies CB resistance.

**Figure 1.**
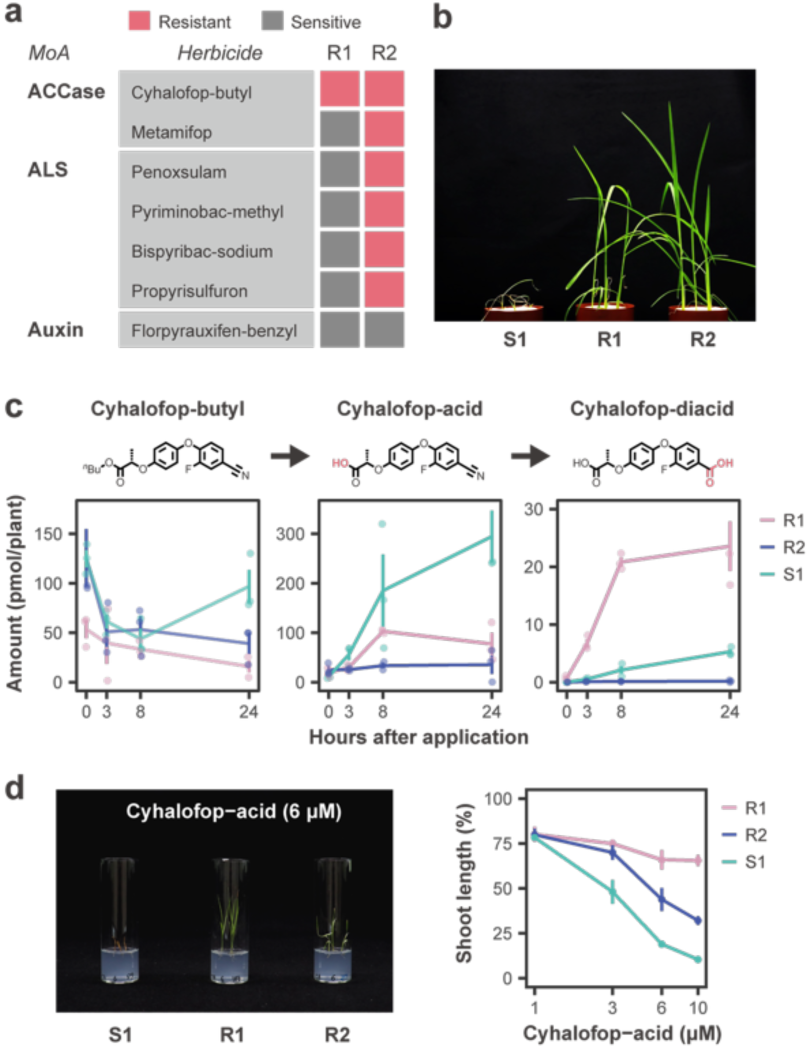
Physiological characterization of cyhalofop-butyl resistance in *Echinochloa crus-galli* a, Previously reported cross-resistance profiles of the susceptible line (S1) and two resistant lines (R1 and R2). **b,** Representative phenotypes of S1, R1 and R2 following foliar application of cyhalofop-butyl. **c,** Time-course LC–MS/MS analysis of cyhalofop-butyl metabolism to cyhalofop-acid and cyhalofop-diacid in S1, R1 and R2. Data represent means ± s.e.m. (n = 3). **d,** Cyhalofop-acid sensitvitiy. Representative images were obtained after culture in liquid medium containing 6 μM cyhalofop-acid. Growth was quantified relative to untreated controls. Data represent means ± s.e.m. (n = 3).

CB is a proherbicide with little direct herbicidal activity and is hydrolysed in plants to cyhalofop-acid, its herbicidally active form.^20^ Further conversion of cyhalofop-acid to cyhalofop-diacid inactivates the herbicide. LC–MS/MS analysis showed that cyhalofop-acid accumulated rapidly in the susceptible S1 line following CB treatment, whereas its abundance remained low in both resistant lines (Fig. 1c). Cyhalofop-diacid accumulated to high levels only in R1, indicating rapid detoxification of cyhalofop-acid in this line. Because CB abundance varied among samples, we directly assessed sensitivity to cyhalofop-acid. Both resistant lines were less sensitive than S1 (Fig. 1d), demonstrating that their resistance extends to the active acid itself. These results identify rapid conversion of cyhalofop-acid to cyhalofop-diacid as the basis of resistance in R1; subsequent analyses therefore focused on this line.

### CB resistance maps to a single major locus on chromosome 7B

To identify the genetic basis of CB resistance in R1, we generated an F2 population by crossing R1 with S1 line (Fig. 2a). Following application of CB at the recommended field rate, all R1 plants survived whereas all S1 plants were killed (Fig. 2b). Among F2 plants, 32 were resistant and 98 were susceptible, fitting a 1:3 segregation ratio (χ^2^ test, *P* = 0.9193).

**Figure 2.**
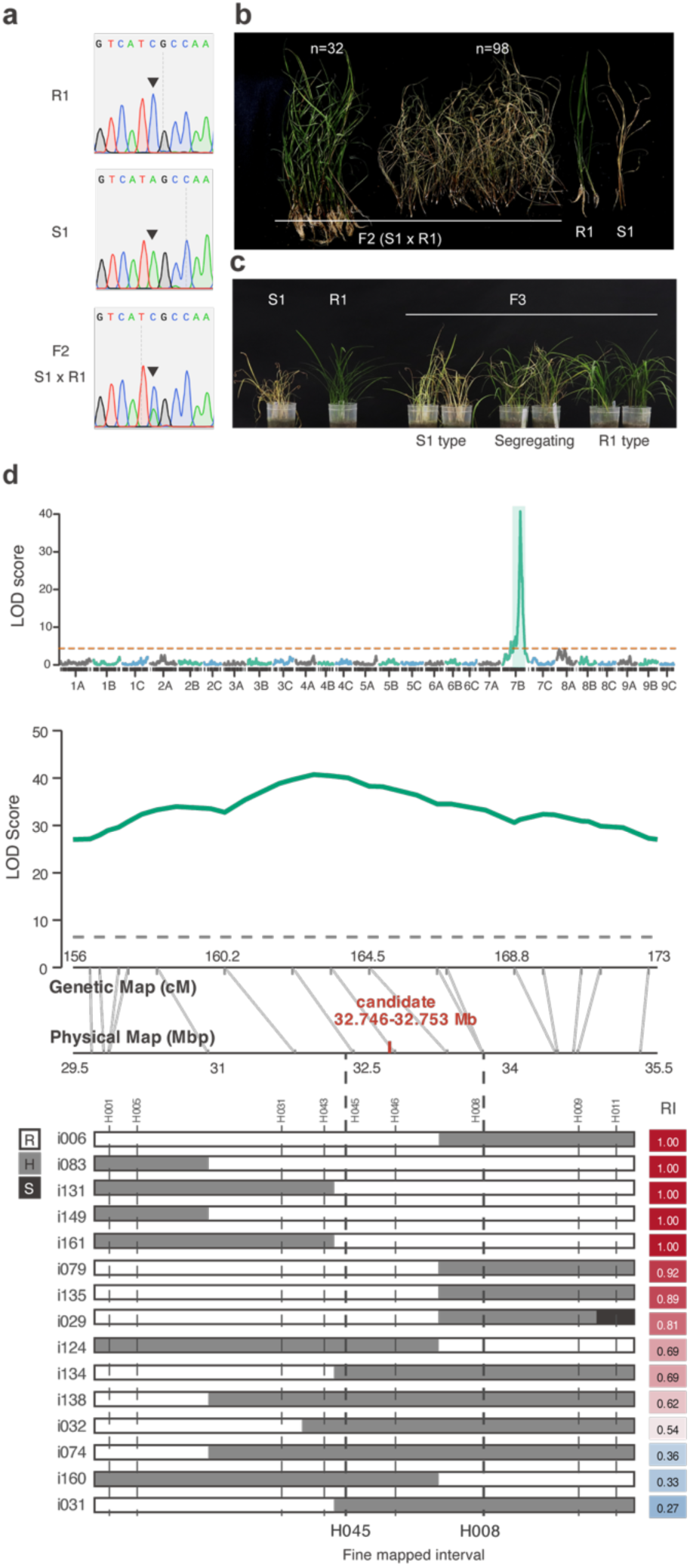
Genetic mapping identifies a major cyhalofop-butyl (CB) resistance locus a, Sequence chromatograms confirming successful hybridization of the S1 × R1 F1 plant at a diagnostic SNP. **b,** Segregation of CB resistance in the F2 population derived from S1 × R1. **c,** Representative phenotypes of F2:3 families derived from resistant and susceptible F2 plants. **d,** Genome-wide LOD profile from interval mapping (IM). The dashed line indicates the significance threshold. **e,** Fine mapping of the resistance locus using recombinant individuals narrowed the candidate interval to a 1.42-Mb region between markers H045 and H008. RI, resistance indices.

For genetic mapping, a separate set of 181 F2 individuals from the same S1 × R1 cross was genotyped by dpMIG-seq,^21^ and the resulting markers were used to construct a linkage map (Fig. S1). QTL analysis was performed using resistance indices derived from the CB responses of F2:3 families generated from 156 of these individuals (Fig. 2c). Interval mapping identified a single major QTL on chromosome 7B, with a peak at 32.80 Mb (LOD = 40.7) explaining 69.9% of the phenotypic variance (Fig. 2d). Composite interval mapping independently identified the same QTL. Fine mapping of 15 recombinant individuals using nine indel markers narrowed the locus to a 1.42-Mb interval between H045 and H008 (32,291,204–33,706,248 bp), containing 24 gene models. Marker H046 at 32.80 Mb showed complete co-segregation with the resistance phenotype.

### *EcNIT4A2* is highly expressed in the R1 line

To identify genes associated with CB resistance in R1, we first compared the leaf transcriptomes of untreated R1, R2 and susceptible S1 plants. A total of 1,372 genes were specifically upregulated in R1 relative to both R2 and S1 (Fig. 3a). We then performed an independent RNA-seq analysis comparing R1 with three CB-susceptible populations collected from geographically distinct regions of Japan. This second analysis identified 304 genes specifically upregulated in R1, 67 of which were shared with the first dataset (Fig. 3b). Of the 24 gene models located within the 1.42-Mb resistance interval (Fig. 2d), only *EcNIT4A2* was consistently upregulated in R1 in both transcriptome analyses. Consistent with its candidacy, RT-qPCR analysis showed that high *EcNIT4A2* expression co-segregated with resistance in S1 × R1 F2:3 families (Fig. 3c).

**Fig. 3.**
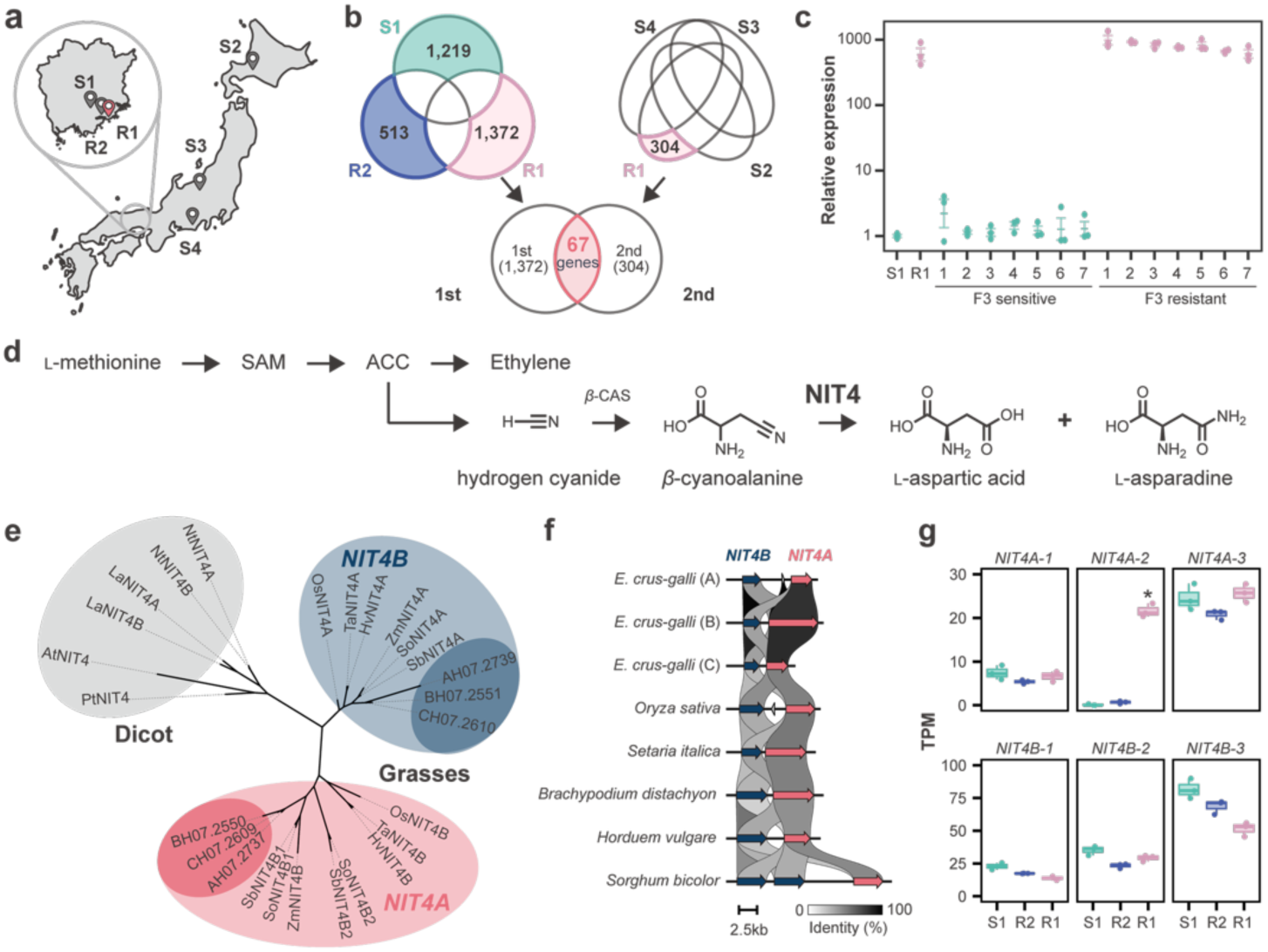
Independent transcriptome analyses identify R1-specific upregulation of *EcNIT4A2*, a nitrilase in the cyanide-detoxification pathway. **a**, Collection sites of the resistant lines R1 and R2 and susceptible lines S1–S4 in Japan. **b,** Venn diagrams of differentially expressed genes identified in two independent RNA-seq analyses. The R1-specific gene sets from the first (R1, R2 and S1) and second (R1 and S2– S4) analyses shared 67 genes. **c,** RT–qPCR analysis of *EcNIT4A2* expression in S1, R1 and seven sensitive and seven resistant F_3_ families. Expression is shown relative to S1. Points represent biological replicates (n=3); bars indicate mean ± s.e.m. **d,** Ethylene biosynthesis and the associated cyanide-detoxification pathway involving β-cyanoalanine synthase (β-CAS) and NIT4. SAM, S-adenosyl-L-methionine; ACC, 1-aminocyclopropane-1-carboxylic acid. **e,** Maximum-likelihood phylogeny of plant NIT4 proteins. **f,** Microsynteny of the *NIT4A* and *NIT4B* regions in grasses. Connecting ribbons indicate sequence identity. **g,** Transcript abundance of the three *EcNIT4A* and three *EcNIT4B* homoeologues in S1, R2 and R1 in the first RNA-seq dataset (n = 3 biological replicates). *, R1-specific increase in *EcNIT4A2* identified by edgeR analysis.

*EcNIT4A2* encodes a NIT4 nitrilase, an enzyme with a conserved function in cyanide detoxification associated with ethylene biosynthesis.^22,23^ Ethylene production generates cyanide as a by-product, which is incorporated into β-cyanoalanine and subsequently hydrolysed by NIT4 to produce asparagine or aspartate (Fig. 3d). Thus, the genetic and transcriptomic analyses identified an enzyme whose endogenous function involves nitrile hydrolysis, the same reaction underlying detoxification of cyhalofop-acid to cyhalofop-diacid.

Whereas NIT4 occurs as a single-copy gene in most dicots, grasses possess two distinct paralogs, NIT4A and NIT4B (Fig. 3e).^24^ These paralogs are typically arranged in tandem in grass genomes (Fig. 3f). In the hexaploid *E. crus-galli* genome, tandem *NIT4A– NIT4B* pairs occur on homoeologous chromosomes 7A, 7B and 7C, resulting in six NIT4 copies. Of these, *EcNIT4A2*, the B-subgenome *NIT4A* copy on chromosome 7B, was uniquely overexpressed in R1 (Fig. 3g).

### *EcNIT4A2* confers herbicide resistance

To determine whether *EcNIT4A2* can confer CB resistance, we ectopically expressed the R1 alleles of *EcNIT4A2* and *EcNIT4B2* in rice under the cauliflower mosaic virus (CaMV) 35S promoter. Whereas *EcNIT4B2* had little effect on CB tolerance, some of the 12 independent *EcNIT4A2*-expressing callus lines survived 300 μM CB, ∼30 times the concentration lethal to wild-type calli (Fig. 4a). *EcNIT4A2* conferred strong CB resistance, whereas *CYP81A68*, *EcGSTU23* and *CYP89B16*—previously reported to confer CB resistance in *E. crus-galli*^25–27^—showed little or no effect under our experimental conditions (Fig. S2).

**Figure 4.**
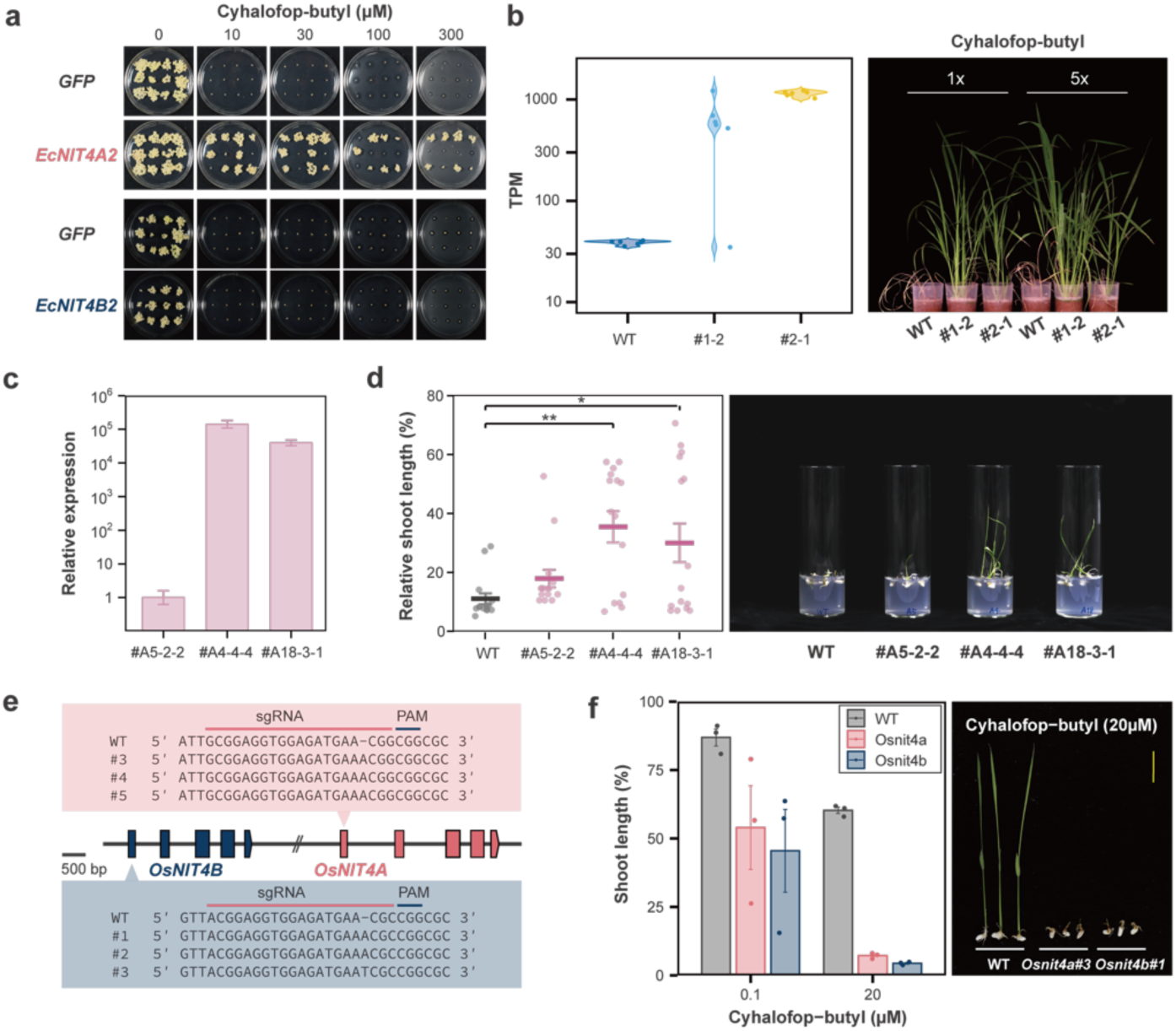
Functional validation of *EcNIT4A2* as the cyhalofop-butyl (CB) resistance gene. a, Rice calli expressing *EcNIT4A2* or *EcNIT4B2* cultured on medium containing CB. **b,** *Echinochloa phyllopogon* plants transformed with *EcNIT4A2*. Relative *NIT4A2* expression, including both transgenic *EcNIT4A2* and endogenous *EpNIT4A2* transcripts, was quantified by RNA-seq. **c,** RT–qPCR analysis of *OsNIT4A* expression in *OsNIT4A*-overexpressing rice plants (*n* = 4). **d,** CB sensitivity was assessed by measuring relative shoot length following treatment with 300 µM CB (n = 15). * and ** indicate *P* < 0.05 and *P* < 0.01, respectively, compared with WT (Dunnett’s multiple-comparison test). Representative phenotypes of rice plants overexpressing *OsNIT4A* following treatment with 300 µM CB. **e,** Design of guide RNAs targeting *OsNIT4A* and *OsNIT4B* in rice. **f,** CB sensitivity of the knockout mutants. Data are presented as mean ± s.e.m (n=3). Yellow bar, 2 cm.

We next expressed *EcNIT4A2* in *E. phyllopogon*, a congeneric rice weed, under the maize ubiquitin promoter using a recently established transformation system.^28^ Two independent transgenic lines were obtained. Both exhibited resistance to CB in callus assays (Fig. S3), and their regenerated progeny were resistant in both foliar application and sterile culture assays (Fig. 4b, Fig. S3). RNA-seq analysis showed that *NIT4A* transcript abundance in resistant transgenic progeny was generally several hundred to over 1,000 TPM, substantially exceeding endogenous *NIT4A2* expression in wild-type *E. phyllopogon* (∼40 TPM). Together, these results demonstrate that overexpression of *EcNIT4A2* is sufficient to confer CB resistance in *Echinochloa*.

We next examined whether overexpression of wild-type rice NIT4A could also enhance CB tolerance. Overexpression of *OsNIT4A* had little effect in callus assays (Fig. S4) but conferred a significant increase in CB tolerance at the whole-plant level (Fig. 4c,d). To assess the contribution of endogenous NIT4 activity to basal CB tolerance, we disrupted *OsNIT4A* and *OsNIT4B* using CRISPR/Cas9. Disruption of either *OsNIT4A* or *OsNIT4B* increased CB sensitivity by approximately two orders of magnitude, as both mutants showed growth inhibition at 0.1 μM CB comparable to or greater than that of WT plants at 20 μM CB (Fig. 4e,f). Grass NIT4A and NIT4B proteins can form heteromeric complexes.^29–32^ If these complexes contribute to cyhalofop-acid detoxification, disruption of either gene would be expected to reduce heteromeric NIT4 activity, potentially explaining why both knockouts showed similar increases in CB sensitivity. Together, these results indicate that endogenous NIT4 activity contributes to basal CB tolerance in rice. Notably, the modest effect of *OsNIT4A* overexpression contrasts with the strong resistance conferred by the R1 *EcNIT4A2* allele.

### D317G enhances NIT4-mediated herbicide detoxification

Comparison of the EcNIT4A2 coding sequences from S1 and R1 identified a single-nucleotide substitution in the R1 allele, resulting in an aspartic acid-to-glycine change at the position corresponding to D317 in AtNIT4 (D315 in EcNIT4A2; Fig. 5a). To facilitate comparison among NIT4A and NIT4B proteins, residue positions are reported using AtNIT4-equivalent numbering throughout; this substitution is therefore referred to hereafter as D317G. This position is well-conserved as aspartic acid (D) among grass NIT4A proteins, including those from rice and *Echinochloa* (Fig. 5b). Functional comparison showed that expression of the D317 *EcNIT4A2* allele from S1 conferred little resistance to CB as observed for rice *NIT4A* (Fig. S4), whereas the G317 allele from R1 conferred strong resistance (Fig. 5c, Fig. S5). Expression of the two other *E. crus-galli NIT4A* homeologues, *EcNIT4A1* and *EcNIT4A3*, both of which encode D317, likewise failed to confer substantial resistance (Fig. 5c). Introduction of D317G into rice NIT4A also resulted in a marked increase in CB resistance (Fig. 5d). Together, these results identify D317G as the coding mutation responsible for enhanced NIT4A-mediated CB resistance in *E. crus-galli*.

**Figure 5.**
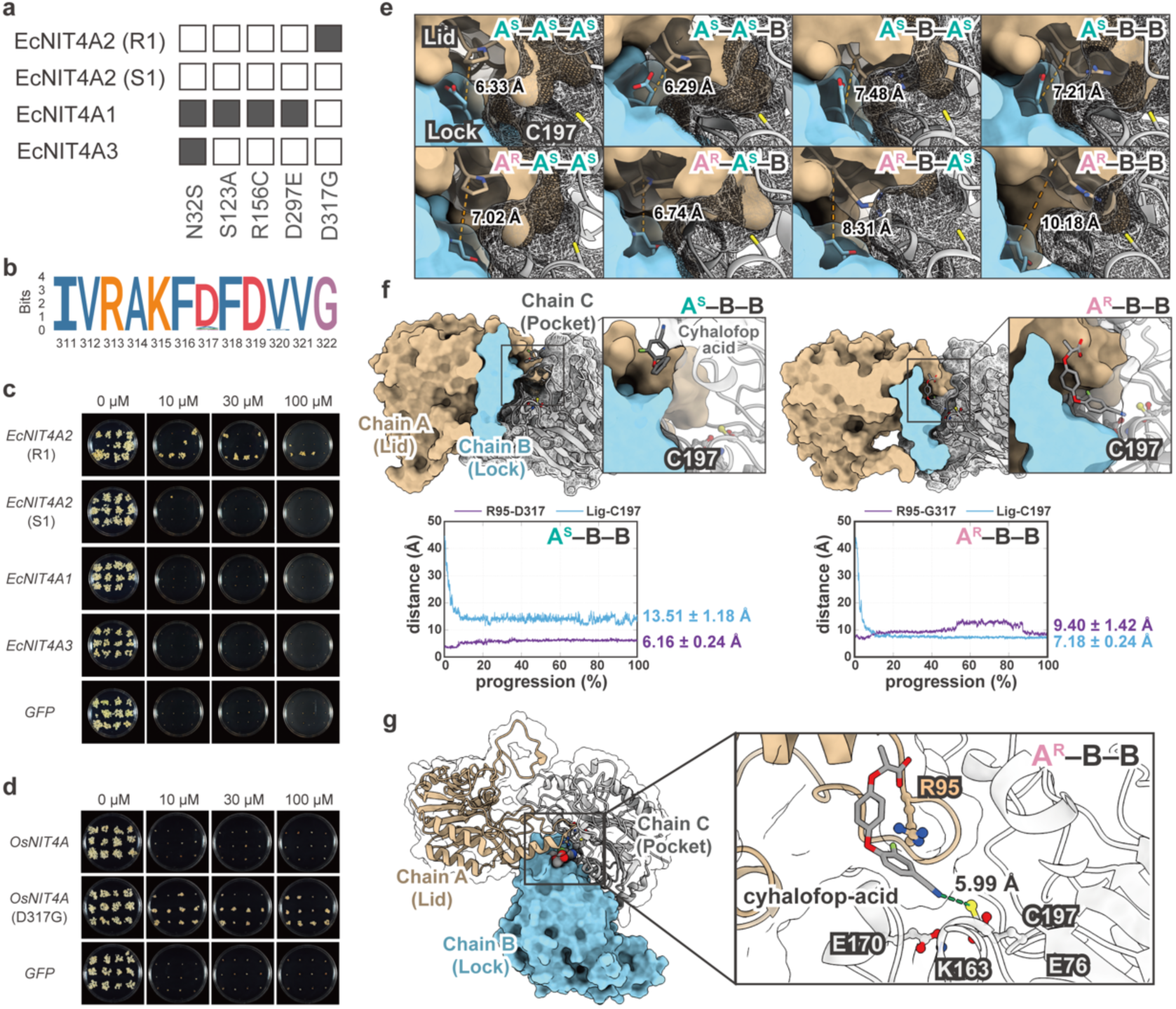
S**t**ructural **basis of enhanced cyhalofop metabolism by the D317G substitution in EcNIT4A2. a,** Distribution of five amino acid substitutions among EcNIT4A2 from R1 and S1 and the homoeologous EcNIT4A1 and EcNIT4A3 proteins. Filled squares indicate residues differing from S1 EcNIT4A2. **b,** Sequence logo of residues 311–322 from grass NIT4A homologues. **c,** Rice calli expressing R1 or S1 *EcNIT4A2*, *EcNIT4A1* or *EcNIT4A3* on medium containing the indicated concentrations of cyhalofop butyl. **d,** Rice calli expressing wild-type *OsNIT4A* or *OsNIT4A* carrying D317G on medium containing the indicated concentrations of cyhalofop butyl. **e,** Eight modelled EcNIT4A–EcNIT4B assemblies (lock–lid–pocket) for the A^S^ (upper panel) and A^R^ (lower panel) variants. Lid–lock distances were measured between residue Cα atoms in the final structures obtained from LB-PaCS-MD simulations. EcNIT4A2 occupies the lock position throughout. **f,** Binding poses of A^S^–B–B (left) and A^R^–B–B (right), showing lock (chain A, tan), lid (chain B, blue) and pocket (chain C, grey). Insets show cyhalofop-acid relative to the catalytic C195. Lower panels, ligand–C195 (blue) and R93–residue 317 (purple) distances over simulation progression (mean ± s.d.). **g,** Minimum-energy conformation of A^R^–B–B. Inset shows cyhalofop-acid packed against R93, with the carboxylate coordinated by K161, E168 and E74 and the nitrile carbon 5.99 Å from C195. Residue positions are reported using AtNIT4-equivalent numbering (PDB 6I5T).

To explain the enhanced activity conferred by D317G, we examined the structural organization of grass NIT4 complexes. The only experimentally determined NIT4 structure is the Arabidopsis cryo-EM complex, in which the substrate pocket is formed by three adjacent subunits (pocket, lid and lock).^33^ In this structure, D317 from the lock subunit forms an intersubunit contact with R95 in the adjacent lid subunit. Arabidopsis has a single NIT4, whereas grasses possess two paralogous subfamilies, NIT4A and NIT4B, proposed to function as heteromers,^29–32^ implying that several NIT4A–NIT4B assemblies may exist. Because residue 317 contributes to the pocket only from the lock subunit^33^ and its potential interacting residue at position 95 differs between the alternative lid subunits (P95 in EcNIT4A2 and R95 in EcNIT4B2), we modelled all eight possible assemblies containing either the S1 or R1 EcNIT4A2 at the lock position and either EcNIT4A2 or EcNIT4B2 at the lid and pocket positions (Fig. 5e).

To compare substrate access among these assemblies, we performed enhanced-sampling molecular dynamics (MD) simulations designed to sample the entry of cyhalofop-acid into the catalytic pocket. Across 200 LB-PaCS-MD cycles, the ligand approached the catalytic pocket in all eight assemblies (Fig. S6a), but the extent of approach varied substantially with subunit composition. A^R^–B–B showed the most favorable substrate access and the widest lid–lock separation among the eight modeled assemblies in the final LB-PaCS-MD structures (10.18 Å; Fig. 5e). Free-energy landscapes supported this trend, with A^R^–B–B reaching the smallest annotated ligand–pocket distance (6.09 Å), compared with 11.55 Å in A^S^–B–B (Fig. S6b). The remaining assemblies fell between these extremes. Together, these results suggest that D317G enhances substrate access in a subunit-dependent manner, with the strongest effect observed in A^R^–B–B.

Extended simulations of A^S^–B–B and A^R^–B–B further supported this difference (Fig. 5f and Supplementary Videos S1 and S2). In A^S^–B–B, the ligand remained near the pocket entrance, with a mean ligand–C197 distance of 13.51 ± 1.18 Å, whereas in A^R^–B–B, it penetrated further into the pocket (7.18 ± 0.24 Å). The R95–residue 317 distance was also greater in A^R^–B–B (9.40 ± 1.42 Å) than in A^S^–B–B (6.16 ± 0.24 Å), consistent with a wider lid–lock opening. Additional contact analyses revealed more persistent ligand–lock interactions in A^S^–B–B and transient R95–D317 hydrogen bonding during the early simulation stages (Fig. S6c,d). In a representative bound conformation of A^R^–B–B (Fig. 5g), the aryl moiety of cyhalofop-acid packs against R95 of the NIT4B lid, while its carboxylate group interacts with E76, K163 and E170, positioning the nitrile carbon 5.99 Å from C197. Together, these findings suggest that D317G facilitates substrate entry by altering the lid– lock geometry and substrate-entry pathway rather than directly modifying the catalytic center.

### A promoter structural variant drives EcNIT4A2 overexpression

To investigate the basis of *EcNIT4A2* overexpression, we compared the genomic region surrounding *EcNIT4A2* between the S1 and R1 genomes assembled using HiFi sequencing. In the S1 genome, *EcNIT4B2* was located upstream of *EcNIT4A2*, with an approximately 4.2-kb intergenic region separating the *EcNIT4B*2 stop codon and the *EcNIT4A*2 start codon (Fig. 6a). The sequence immediately upstream of the *EcNIT4A2* coding region was identical between S1 and R1 for the first 106 bp. However, the adjacent 3,349-bp region in S1 was replaced by a 313-bp sequence in R1 (Fig. 6a). The remaining intergenic sequence was nearly identical between the two alleles.

**Figure 6.**
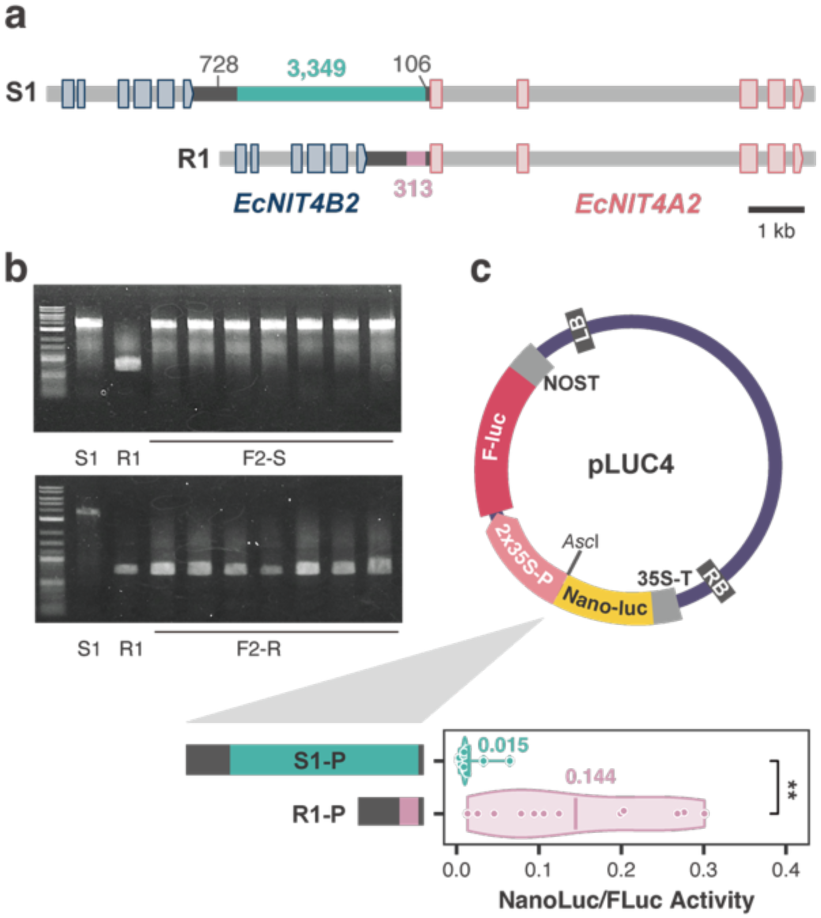
A promoter structural variant enhances EcNIT4A2 expression a, Comparison of the *EcNIT4A2* promoter regions in S1 and R1. **b,** PCR validation of the promoter structural variant in representative susceptible and resistant F_3_ plants. **c,** Promoter activity of the S1 and R1 *EcNIT4A2* promoter alleles measured using a dual-luciferase reporter assay in rice calli. Firefly luciferase activity was normalized to NanoLuc luciferase. **, *P* < 0.01 (two-sided Student’s *t*-test).

To further examine the relationship between promoter genotype and *EcNIT4A2* expression, we genotyped the promoter region in the progeny previously analyzed by qRT-PCR (Fig. 3g). All individuals carrying the R-type promoter exhibited elevated *EcNIT4A2* expression, whereas individuals carrying the S-type promoter showed expression levels comparable to the susceptible parent (Fig. 6b), indicating complete co-segregation of the promoter SV and *EcNIT4A2* overexpression.

To test whether this promoter SV contributes to *EcNIT4A2* overexpression, the *EcNIT4B2*–*EcNIT4A2* intergenic regions from the S1 and R1 lines were fused to a NanoLuc reporter and analyzed using a dual-luciferase assay in transiently transformed rice calli (Fig. 6c). Whereas the S1 promoter drove only weak reporter expression, the R1 promoter exhibited approximately 10-fold higher normalized NanoLuc activity (NanoLuc/F-LUC). Together, these findings demonstrate that a promoter SV is responsible for *EcNIT4A2* overexpression, thereby providing the regulatory basis for its recruitment into herbicide detoxification.

## Discussion

The co-option of pre-existing biological functions is an important mechanism of evolutionary innovation, yet how such transitions occur over contemporary evolutionary timescales remains poorly understood. Here, we demonstrate that resistance to the herbicide cyhalofop-butyl evolved through the co-option of EcNIT4A2, a nitrilase involved in ethylene-associated cyanide metabolism. Resistance arose through the combined effects of a promoter structural variant that markedly increased *EcNIT4A2* expression and a coding mutation that enhanced its herbicide-metabolizing activity. In contrast, the independently resistant R2 line showed neither elevated *EcNIT4A2* expression nor cyhalofop-diacid accumulation, suggesting that alternative resistance mechanisms may operate in this line. Together, our findings provide a molecular example of how regulatory and coding changes can jointly recruit and optimize a conserved metabolic enzyme during rapid adaptation under anthropogenic selection.

The evolutionary history of plant nitrilases suggests that this enzyme family is particularly amenable to functional diversification.^23^ Although ancestral NIT4 enzymes function in cyanide detoxification, gene duplication has repeatedly been followed by changes in substrate specificity. In Brassicaceae, for example, duplicated NIT1–3 enzymes acquired the ability to metabolize glucosinolate-derived nitriles,^24,33,34^ whereas in sorghum, duplication and functional divergence of *NIT4B* gave rise to paralogs specialized for dhurrin turnover.^29^

The evolution of cyhalofop metabolism by EcNIT4A2 extends this recurring pattern of nitrilase diversification into the context of contemporary adaptation. Unlike the endogenous metabolites involved in these earlier examples, CB is a synthetic herbicide introduced only in the 1990s,^35^ illustrating how endogenous metabolic enzymes can be recruited to detoxify entirely novel anthropogenic compounds over short evolutionary timescales.

Resistance in R1 involved changes in both *EcNIT4A2* expression and the function of its encoded protein. Elevated expression of detoxification genes is common in metabolic herbicide resistance,^e.g. 36–38^ whereas direct evidence that adaptive amino acid substitutions can enhance herbicide-metabolizing activity remains scare. In the R1 line, a promoter SV markedly increased *EcNIT4A2* expression, while the D317G substitution substantially enhanced its capacity to metabolize cyhalofop-acid. Importantly, disruption of either *OsNIT4A* or *OsNIT4B* greatly increased CB sensitivity, indicating that endogenous NIT4 complexes already contribute to basal cyhalofop-acid detoxification in plants. The Arabidopsis NIT4 structure revealed an R95–D317 lid–lock interaction, and D317 was proposed as a target for altering substrate size.^33^ Our structural analyses suggest that the naturally occurring D317G substitution disrupts this interaction, widening the substrate-entry channel without altering the catalytic tetrad. This effect was observed only in heteromeric assemblies, consistent with the proposed heteromeric organization of grass NIT4.^29–32^ Together, these findings suggest that herbicide resistance arose through enhancement of a pre-existing metabolic capacity rather than through the emergence of an entirely new herbicide-metabolizing activity.

The evolutionary origins of the regulatory and coding changes underlying EcNIT4A2-mediated resistance remain unresolved. One explanation for the rapid evolution frequently observed in metabolic herbicide resistance is that selection acts primarily on standing genetic variation rather than relying solely on newly arising beneficial mutations.^39^ Notably, a promoter configuration resembling the R1 allele is present in the published genome of an *E. crus-galli* accession originating from China,^40^ raising the possibility that at least the regulatory component of resistance arose from pre-existing variation. In contrast, D317 is highly conserved among plant NIT4 proteins, suggesting stronger evolutionary constraint on this coding position. Determining whether the regulatory and coding variants arose independently, sequentially or from pre-existing variation will require broader population-level analyses of *EcNIT4A2* diversity.

Because NIT4 performs an important physiological role in cyanide detoxification, its recruitment for herbicide metabolism raises the question of how such evolutionary modification could occur without compromising its endogenous function. One possible explanation is functional redundancy. *E. crus-galli* is hexaploid and carries multiple NIT4 homeologues, and *EcNIT4A2* is expressed at only very low levels in the susceptible lines (Fig. 3g). Thus, this particular copy may make only a limited contribution to basal NIT4 function, potentially allowing it to diverge while other copies maintain the ancestral function. Gene duplication is widely recognized as a mechanism that facilitates such evolutionary innovation by relaxing functional constraints on individual copies.^41,42^ Consistent with this interpretation, resistant plants carrying the highly expressed D317G allele showed no obvious developmental defects under our experimental conditions, although the effect of D317G on native β-cyanoalanine metabolism remains unresolved. Functional redundancy among *NIT4* copies may therefore have provided the evolutionary flexibility that enabled *EcNIT4A2* to be co-opted for herbicide detoxification.

Repeated recruitment of conserved genes and pathways is a common feature of evolutionary innovation over deep evolutionary timescales.^3^ Our findings show that the same principle can operate during contemporary adaptation, with regulatory and coding changes jointly enhancing a pre-existing metabolic capacity within only a few decades. More broadly, this work provides a molecular example of how conserved metabolic functions can be co-opted and optimized under intense anthropogenic selection.

## Methods

### Plant materials

*Echinochloa crus-galli* var. *formosensis* lines S1 (formerly Ecf99), R1 (formerly Ecf108), and R2 (formerly Ecf27) were used. The lines were originated in Okayama city, Japan. The details of herbicide sensitivity are described in elsewhere.^18,19^ The crossing of S1 and R1 was performed following hot water emasculation method in rice.^43^ The success of hybridization was confirmed by sequencing the *ALS3* gene in the resultant plants, following PCR amplification with primers listed in Table S1. The F1 plant was self-pollinated to generate F2 progeny. Individual F2 plants were subsequently self-pollinated to produce F2:3 families.

*E. crus-galli* var. *crus-galli* lines EcTU018, EcTU019, and EcTU006, originating from Hokkaido, Niigata, and Aichi, respectively, were designated S2, S3, and S4, respectively. They were all sensitive to CB (Fig. S7). *E. phyllopogon* AEC02-17 line^28^ and rice (*Oryza sativa* L. ‘Nipponbare’) were used for transformation.

### CB sensitivity assay

Foliar application of a commercial formulation of CB was performed essentially as described previously.^19^ Briefly, plants were grown under controlled environmental conditions (28 ± 5 °C, 13-h photoperiod). At the 4- or 5-leaf stage, plants were treated with Clincher 30% EW (Corteva Japan) supplemented with 0.1% (v/v) Surfactant 30 (Maruwa Biochemical) at 300 g active ingredient (a.i.)/ha. Herbicide sensitivity was evaluated 10 or 21 days after treatment, depending on the experiment.

For sensitivity assays in culture tubes, seeds were surface-sterilized by vacuum infiltration in sodium hypochlorite (∼2.5% active chlorine) containing 0.01% (v/v) Tween 20 for 15 min and then washed three times with sterile water. The sterilized seeds were germinated under 25°C and transferred to MS medium supplemented with 0.05% (v/v) Plant Preservative Mixture (Plant Cell Technology). *E. crus-galli* seedlings were grown at 25 °C for 7 days, whereas rice seedlings were grown at 30°C for 5 days, both under a 12-h photoperiod.

For rice callus sensitivity assays, 9–12 independent transgenic calli were cultured on N6D medium containing CB, as described previously. ^37^ For *E. phyllopogon* assays, two independently transformed callus lines were examined, with 12 callus pieces per line assayed on a single plate at each CB concentration.

### Vector construction

Unless otherwise indicated, coding sequences were PCR-amplified from cDNA using the primers listed in Table S1. The PCR products were subcloned into the pCR-Blunt vector using the Zero Blunt PCR Cloning Kit (Thermo Fisher Scientific). The coding sequences of *CYP81A68* (OK483200.1) and *EcGSTU23* (EC_v6.g055562) were synthesized by GenScript (Tokyo, Japan). The coding sequence of *CYP89B16* (Eph1.1ch9A.g032540.1), previously identified in the R511 line of *E. phyllopogon*,^44^ was amplified from R511 cDNA. The D317G substitution was introduced into the *OsNIT4A* coding sequence by site-directed mutagenesis using the primers listed in Table S1.

For construction of overexpression vectors, pCAMBIA1390-sGFP^45^ was digested with BsrGI and SalI, and the respective coding sequences were inserted into the linearized vector using either the In-Fusion HD Cloning Kit (Takara) or the SLiCE method.^46^ The pAHC25^47^ vector was digested with SmaI and SacI, and the *EcNIT4A2* coding sequence was inserted into the linearized vector using SLiCE. Binary CRISPR/Cas9 vectors carrying gene-specific single-guide RNAs were constructed as described previously.^48^ The protospacer– PAM sequences used for *OsNIT4A* (5′-GCGGAGGTGGAGATGAACGGCGG-3′) and *OsNIT4B* (5′-ACGGAGGTGGAGATGAACGCCGG-3′) were designed using CRISPR-P v2.0.^49^

To construct the dual-luciferase reporter vectors, the firefly luciferase (FLUC) coding sequence together with the NOS terminator (NOS-T) was PCR-amplified from pGWB535 (Addgene plasmid no. 74873) using primer set pL1. The 2×CaMV 35S promoter was amplified from pCAMBIA1390-sGFP using primer set pL2. These fragments were assembled into the linearized pUC19 vector supplied with the In-Fusion HD Cloning Kit to generate a 2×35S::FLUC::NOS-T construct. The resulting 2×35S::FLUC::NOS-T cassette was subsequently amplified using primer set pL3. Separately, the 35S terminator (35S-T) was amplified from pB2GW7^50^ using primer set pL4, and the NanoLuc luciferase (NLUC) coding sequence was amplified from pNL1.1.PGK (Promega) using primer set pL5. These fragments were assembled into the linearized pUC19 vector supplied with the In-Fusion HD Cloning Kit to generate an intermediate construct. The intermediate plasmid was digested with AscI, and the 2×35S::FLUC::NOS-T cassette was inserted by In-Fusion cloning to generate pLUC2.

To construct pLUC4, pB2GW7 was digested with MauBI and HindIII. The NOS-T::FLUC::2×35S::NLUC::35S-T cassette was PCR-amplified from pLUC2 using primer set pL6 and inserted into the linearized pB2GW7 backbone by In-Fusion cloning. The S1 and R1 *EcNIT4A2* promoter fragments were PCR-amplified from genomic DNA and subcloned into the pCR-Blunt vector. The promoter fragments were re-amplified using primer set pL7 and inserted into AscI-linearized pLUC4 by In-Fusion cloning. All final constructs were verified by Sanger sequencing. All primers used for vector construction are listed in Table S1.

### Plant transformation

Rice transformation was performed as previously described.^37^ Genome-edited T0 plants were genotyped by Sanger sequencing of PCR amplicons spanning the target site (Table S1), and biallelic mutants were selected for further analyses.

*E. phyllopogon* transformation was performed as previously described.^28^ Briefly, embryogenic calli were induced on MS medium^51^ supplemented with 2.0 mg L^-1^ 2,4-D, 0.1 mg L^-1^ 6-benzylaminopurine and 5 µM copper sulfate. The embryogenic calli were transformed using a self-built particle bombardment apparatus.^52^ Bombardment was carried out using gold particles (1.0 µm diameter; Biorad). The transformants were selected with 5 mg L^-1^ bialaphos. Plants were regenerated by transferring the calli on MS medium supplemented with 0.02 mg L^-1^ naphthaleneacetic acid (NAA) and 2 mg L^-1^ kinetin and 5 mg L^-1^ bialaphos. Regenerated plants were tested for the presence of the EcNIT4A2-NOS terminator by PCR (Table S1). Subsequently, PCR-positive plants were transferred to soil and grown to maturity.

### Dual luciferase assay

Calli co-cultured with Agrobacterium for 1 day at 28°C were washed with sterile water, and 50 mg of callus tissue was snap-frozen in liquid nitrogen and stored at −80°C until use. The frozen tissue was ground and resuspended in Passive Lysis Buffer (Promega). Firefly and NanoLuc luciferase activities were measured using the Nano-Glo Dual-Luciferase Reporter Assay System (Promega) according to the manufacturer’s instructions. Luminescence was detected using a GloMax Explorer (Promega).

### Real-time PCR

Total RNA was extracted from the shoot of a single plant with three biological replicates, and 500 ng of RNA was used to synthesize cDNA with the PrimeScript IV 1st strand cDNA Synthesis Mix (Takara). Quantitative PCR was performed using THUNDERBIRD Next SYBR qPCR Mix (TOYOBO) on a QuantStudio 3 (ThermoFischer). The primers used in this study are listed in Table S1. Primer concentrations for *EcEIF4B* and *EcNIT4A2* were 100 μM and 300 μM, respectively. Samples with *EcNIT4A2* signals below the detection limit after 40 cycles were assigned a Ct value of 40 for downstream analyses. Data was analyzed using the ΔΔCt method.^53^

### Nucleic acid extraction

Total RNA was extracted using a Plant Total RNA Mini Kit (Favorgen). The extracted RNA was treated with TURBO DNA-*free* Kit (Thermo Fisher Scientific). For PCR and dpMIG-seq analysis, DNA was extracted from leaves of each plant with the cetyltrimethylammonium bromide method.^54^ For HiFi sequencing, DNA was extracted using NucleoBond HMW DNA (Takara).

### Genome assembly

HMW DNA was sequenced on a PacBio Revio platform at BGI using their standard library preparation protocol. HiFi reads were assembled with hifiasm (v0.25.0).^55^ Nuclear contigs were extracted using Tiara (v1.0.3)^56^ and aligned to the reference genome of *E. crus-galli*^40^ using RagTag (v2.1.0).^57^ Gene annotation was transferred from the reference genome using Liftoff (v1.6.3).^58^

### RNA-sequencing analyses

Germinated plants (R1, R2, and S1) were grown hydroponically to the 2.5-leaf stage in a growth chamber at 25°C under a 12-h photoperiod (∼300 μmol m⁻² s⁻¹) as described previously.^59^ Shoots from three plants were snap-frozen in liquid nitrogen and stored at −80 °C until RNA extraction. RNA samples were sent to AZENTA for library preparation using their standard protocol. The libraries were sequenced on the HiSeq 4000 platform (150-bp paired-end reads).

For the other RNA-seq experiments, RNA was extracted from the shoots of 2-leaf-stage plants grown in a growth chamber under a 14-h photoperiod at 25 ± 5°C with a high-pressure sodium lamp. Libraries for S1, S2, S3, and S4 were prepared using the NEBNext Ultra II RNA Library Prep Kit for Illumina (NEB, Tokyo, Japan). Sequencing was performed by AZENTA on a NovaSeq 6000 platform (150-bp paired-end reads). RNA samples from transgenic *E. phyllopogon* were sent to BGI for library preparation using their standard protocol. The libraries were sequenced on the DNBSEQ platform (150-bp paired-end reads).

Except for the clean data from BGI, raw data were processed using fastp (v0.23.2)^60^ with the following parameters: -q 30 -l 120 -w 16 --detect_adapter_for_pe. Filtered reads were mapped to the reference genome using STAR (v2.7.9),^61^ and read counts were quantified using RSEM (v1.3.2).^62^ Differentially expressed genes were identified using the R package edgeR (v3.38.4).^63^ Genes specifically upregulated in each line were identified by comparing the target line with the other lines using pairwise contrasts in a generalized linear model framework. Genes were defined as specifically upregulated when they showed significant upregulation in both comparisons (FDR < 0.05 and logFC > 1).

### dpMIG-seq and genetic analysis

Genomic DNA was extracted from F2 individuals as well as the parental lines (S1 and R1). dpMIG-seq libraries were prepared as reported previously.^21^ Paired-end sequencing (150 bp) were performed on a DNBSEQ-G400 platform. Raw reads were processed using fastp (v0.23.4)^60^ with the parameters -q 30 -l 120 -w 16 --detect_adapter_for_pe. Filtered reads were aligned to the *E. crus-galli* genome using strobealign (v0.15.0),^64^ and alignments were coordinate-sorted with SAMtools (v1.11).^65^ BAM files were filtered using bamutils (-mismatch 3 -properpair) (v0.5.9).^66^ Variants were called using SAMtools mpileup and BCFtools BCFtools (v1.19).^65^

A linkage map was constructed using LEP-MAP3 (Rastas, 2017).^67^ Markers were assigned to chromosomes based on their physical positions in the reference genome.^40^ Markers showing segregation distortion or linkage relationships inconsistent with the reference genome were excluded, and marker order and genetic distances were estimated using the Haldane mapping function.

QTL analysis was performed using the R/qtl package (Broman et al., 2003).^68^ The F_2_ individuals were genotyped by dpMIG-seq, and resistance phenotypes of the corresponding F_2:3_ families were used for QTL mapping. Interval mapping and composite interval mapping were performed, and genome-wide significance thresholds were determined by 1,000 permutation tests. Confidence intervals were defined as 1.5-LOD support intervals. Additive and dominance effects, together with the proportion of phenotypic variance explained, were estimated for the detected QTL.

To refine the resistance locus, recombinant individuals identified within the QTL region by dpMIG-seq were selected for fine mapping. Nine indel markers were developed from whole-genome resequencing data of the QTL region (Table S1) and used to genotype the recombinant individuals. The QTL interval was refined by comparing graphical genotypes with resistance phenotypes and identifying recombination breakpoints delimiting the resistance locus.

### LC-MS/MS

4-leaf stage plants were treated with 5 mM CB by applying 1 μL at each of three spots (3 μL per plant). Plants were sampled at 0, 3, 8, and 24 h after application, and two plants were pooled as one sample. Each sample was extracted with 300 μL of 80% (v/v) methanol and transferred to a 1.5-mL microcentrifuge tube. After centrifugation at 15,000 rpm. for 10 min, the supernatant was subjected to LC–MS/MS analysis. Metabolites were analyzed using a Nexera X2 UHPLC system (Shimadzu) coupled to a TripleTOF^TM^ 5600+ mass spectrometer (SCIEX) equipped with an electrospray ionization (ESI) source (DuoSpray^TM^ ion source). Chromatographic separation was performed on a SunFire C18 column (2.1 × 150 mm, 3.5 μm; Waters) fitted with an OptiGuard C18 guard column (1.0 × 15 mm, Optimize Technologies) maintained at 40°C. The mobile phases consisted of (A) water containing 10 mM formic acid and 10 mM ammonium formate and (B) methanol. The gradient was 10% B at 0 min, increased linearly to 95% B at 15 min, maintained until 25 min. The injection volume was 1 μL.

CB was analyzed in positive ion mode, whereas cyhalofop-acid and cyhalofop-diacid were analyzed in negative ion mode. The ion spray voltages were +5500 V and − 4500 V for positive and negative modes, respectively, and the source temperature was 550°C. CB, cyhalofop-acid, and cyhalofop-diacid were quantified by targeted LC–MS/MS using the transitions *m/z* 375.1 ([M+NH4]+) → 256.07793, *m/z* 300.1 ([M-H]^−^) → 228.04663, and *m/z* 319.1 ([M-H] ^−^) → 247.04121, respectively.

### Phylogeny analyses of NIT4 genes

Amino acid sequences of NIT4 of *A. thaliana*, *Nicotiana tabacum*, *Lupinus angustifolius*, *Saccharum officinarum*, *Hordeum vulgare*, *O. sativa*, *Populus trichocarpa*, *Triticum aestivum* and *Zea mays* were aligned with MAFFT (v7.520).^69^ A maximum-likelihood phylogenetic tree was inferred from the aligned sequences using IQ-TREE (v.3.1.2)^70^ and visualized using iTOL (v7.6).^71^ The local synteny around NIT4 was visualized with clinker (v0.0.21).^72^ Sequence logo was generated with grass NIT4A sequences (Table S2) using ggseqlogo (v0.2).^73^

### Homology Modeling and MD simulation of NIT4 fibril

Eight distinct NIT4 fibril systems were constructed from trimeric assemblies where each monomer contributed specific pocket, lid, and lock regions. The pocket and lid regions were assigned to either EcNIT4A2_S (designated as S) or EcNIT4B2 (designated as B), whereas the lock region was assigned to either EcNIT4A2_S (S) or EcNIT4A2_R1 (designated as R). This combinatorial scheme yielded eight systems designated as A^S^–A^S^–A^S^, A^S^–B–B, A^S^–A^S^– B, A^S^–B– A^S^, A^R^–A^S^–A^S^, A^R^–B–B, A^R^–A^S^–B, and A^R^–B– A^S^, where the three-letter code indicates the identity of the pocket, lid, and lock regions, respectively. Structural models were generated using SWISS-MODEL^74^ with cryo-EM structure 6I5T^33^ as the template. Each modeled trimeric fibril complex was prepared for MD simulation using the AMBER24 package.^75^ Protein topologies were generated with the AMBER ff19SB force field, while parameters for cyhalofop-acid ligand were derived from the General AMBER Force Field. Ligand atomic partial charges were assigned via the restrained electrostatic potential method following density functional theory calculations at the B3LYP/6-31G(d) level using Gaussian16.^76^ The ANTECHAMBER module was employed to generate ligand topology and parameter files compatible with the AMBER force field. Complexes were solvated in truncated octahedral TIP3P boxes with a 10 Å buffer and neutralized with Na^+^ or Cl^−^. Systems were minimized for 1000 restrained steepest-descent and 1000 unrestrained conjugate-gradient steps, heated from 0 to 300 K over 100 ps under NVT, and equilibrated for 1 ns under NPT at 300 K and 1.013 bar. Langevin dynamics (2.0 ps^−1^) and the Monte Carlo barostat were used. All MD simulations employed a 2 fs integration time step with the SHAKE algorithm applied to constrain bonds involving hydrogen atoms. Long-range electrostatic interactions were treated with the particle mesh Ewald method using a real-space cutoff of 10 Å, and the same cutoff was applied for short-range van der Waals interactions.

### Enhanced Sampling via LB-PaCS-MD Simulation

To efficiently sample the binding pathway of cyhalofop-acid into the fibril pocket, the Ligand-Binding Parallel Cascade Selection Molecular Dynamics (LB-PaCS-MD) approach was employed through the PaCS-Q workflow.^77^ For each of the eight systems, enhanced sampling was conducted over 220 cycles, with 5 independent replicas (candidates) per cycle and a production length of 100 ps per replica. At the end of each cycle, candidate trajectories were ranked based on the ligand–pocket center-of-mass distance (*d*_Lig–pocket_), and the five snapshots exhibiting the shortest distances were selected as seed structures for the next cycle. The selection algorithm employed a likelihood-maximization approach with additional weighting factors to prevent artificial biasing toward unrealistic conformations. Selection parameters were optimized through preliminary validation runs to ensure adequate sampling of intermediate states. This iterative selection scheme progressively biases the ensemble toward the bound state while preserving the stochastic nature of unbiased MD within each short replica. Convergence was assessed by monitoring the evolution of *d*_Lig–pocket_ and ensuring that fluctuations in this distance reached a plateau over the final 50 cycles.

### Free-Energy Landscape and Binding-State Analysis

Two-dimensional free energy landscapes (2D-FEL) were constructed using *d*_Lig–pocket_ and the number of protein-ligand contacts as reaction coordinates. Contact analysis was performed using a distance cutoff of 4.0 Å between heavy atoms. Free energy surfaces were calculated using the pyEMMA^78^ package with Markov State Model (MSM) analysis, discretizing data into 50×50 grids and computing free energy as F(i,j) = -*k*_B_*T* ln P(i,j), where P(i,j) represents the probability of observing the system in bin (i,j). Global minimum states (GMS) were identified as the lowest free energy regions in each 2D-FEL. Statistical significance of binding affinity differences was assessed through bootstrapping analysis of trajectory data (n=1000 bootstrap samples). Representative structures from GMS were extracted and analyzed for detailed protein-ligand interactions using VMD and in-house Python scripts.

### Statistics

For box plots, the centre line indicates the median, boxes represent the interquartile range (IQR), whiskers extend to 1.5 × IQR, and points indicate individual biological replicates. Violin plots show the kernel density distribution. The centre line indicates the median, boxes indicate the interquartile range, whiskers extend to 1.5 × IQR, and points represent individual biological replicates. Two-sided *t*-test was permed using the tidyplots R package.^79^ Relative shoot height was analyzed by one-way ANOVA followed by Dunnett’s multiple-comparison test comparing each transgenic line with WT, using the multcomp R package.^80^

## Acknowledgements

We thank Dr. Hiroyuki Kasahara for helpful discussions on the enzymatic properties of NIT4. Computations were partially performed on the NIG supercomputer at ROIS National Institute of Genetics. This work was supported in part by the JSPS KAKENHI Grant (JP19K06066, to A.U., JP23K23612, to S.I.), JST PRESTO (JPMJPR24N2, to S.I.), Japan Association for Advancement of Phyto-Regulators (JAPR) (JA-2022-1 to A.U.), and the FLOuRISH Fellowship Program of TUAT to D.S.

## Author contribution

S.I. and A.U. conceived the project. H.I., L.N., S.I. and A.U. characterized plants. M.I. and C.A.T conducted herbicide metabolism study. L.N., D.S. and S.I. designed and performed dual luciferase assay. H.I. and S.I. conducted RNA-seq analyses. L.N., S.I. and M.P.S. conducted genome analyses. H.I., L.N., M.S., T.G. and S.I. transformed plants. M.E. generated rice knockout mutants. K.Y. provided *E. phyllopogon* seeds. K.N. and H.I. performed genetic mapping. K.H., Y.S. and R.S. performed structure analysis. S.I., H.I., L.N. and D.S. performed other data analyses. S.I., K.H. and K.N. wrote the paper with feedback from all authors.

## Ethics declarations

Competing interests

The authors declare no competing interests.

