## Supplemental data for "Evolution of herbicide resistance through co-option of an ethylene-associated nitrilase"

### Supplemental Table 1. Primers used in this study.

| Purpuse | Gene/primer set | Gene ID | Forward primer (5'-3') | Reverse primer (5'-3') |
|---|---|---|---|---|
| Subcloning | *EcNIT4A1* | AH07.2739 | CAGCGTCCACTCCCAGTC | GTCCCAGAAGATCCAACAGGA |
| | *EcNIT4A2* | BH07.2551 | CGTTGCTCAGCATCCACA | GTGCCAGGTTCATCAACGTATAC |
| | *EcNIT4A3* | CH07.2610 | CCCAAGCAAGTGGCAGTAGTA | CCGATGCCTACGCTGGTC |
| | *EcNIT4B2* | BH07.2550 | TCCACGGTCTCCCTCTTCA | GCCATGATCCGTTACCACCT |
| | *OsNIT4A* | Os02g0635200 | TCCCGCGTCCACTTCGAG | AATGATTCACCGTCTAGTGACAG |
| | *OsNIT4B* | Os02g0635000 | TTCCCGCTCAATTTCTTGG | ATTTTATTGCACACCGCGA |
| pCAMBIA vector construction | *EcNIT4A1* | AH07.2739 | TACAATTACAGTCGAATGGCTCTGGTGACCTCG | CCGCTTTACTTGTACTTAGTAGGACCTAGCAGTACTATCA |
| | *EcNIT4A2* | BH07.2551 | TACAATTACAGTCGAATGGCTCTGGTGACCTCG | CCGCTTTACTTGTACTCAGTAGGACCTAGCAGTACTATCA |
| | *EcNIT4A3* | CH07.2610 | TACAATTACAGTCGAATGGCTCTGGTGACCTCG | CCGCTTTACTTGTACTCAGTAGGACCTAGCAGTACTATCA |
| | *EcNIT4B2* | BH07.2550 | TACAATTACAGTCGAATGTCCCTGGCTCCAGCG | CCGCTTTACTTGTACTTAGGTCTTCAAAACGTCATC |
| | *OsNIT4A* | Os02g0635200 | TACAATTACAGTCGAATGGCTATGGTCCCCTC | CCGCTTTACTTGTACTTAGTAGGGCTTTGCAGTG |
| | *OsNIT4B* | Os02g0635000 | TACAATTACAGTCGAATGGCTTTGGCCCCCG | CCGCTTTACTTGTACTTAATGGCCATCAGGTCCT |
| | *CYP81A68* | OK483200.1 | TACAATTACAGTCGAATGGATAAGGCCTACATCGC | CCGCTTTACTTGTACTCAGAGCTCCTGAAGCACAC |
| | *EcGSTU23* | EC_v6.g055562 | TACAATTACAGTCGAATGGCAGACAAGGGCGTG | CCGCTTTACTTGTACTTACTGCGCGGAGAGCAG |
| pAHC25 vector construction | *EcNIT4A2* | | CTTCTGCAGGTCGACATGGCTCTGGTGACCTCG | GATCGGGGAAATTCGTCAGTAGGACCTAGCAGTACTATCAC |
| Site directed mutagenesis | *OsNIT4A* | | TGCGAGCCAAGTTTGGTTTTGATGTGGTGGG | CCCACCACATCAAAACCAAACTTGGCTCGCA |
| Real-time PCR | *EcNIT4A2* | | CCCTGCCCTAGATATCCTTAAC | GACCGAGATAGTACAAACCCCTAT |
| | *EcEIF4B* | | GGCCAGAGTCAGATGAAGAAA | GAGGCTTGGTCAGAACCATC |
| pLUC construction | pL1 | | ATCTATCTCTATGGAAGACGCCAAAAACATAAAG | CAGGTCGACTCTAGAGGATCCGCGCGCGGATCTAGTAACATAGATGACACCGC |
| | pL2 | | GAGCTCGGTACCCGGGGATCATGGTGGAGCACGACACT | CGTCTTCCATAGAGATAGATTTGTAGAGAGAGACT |
| | pL3 | | GAGCTCGGTACCCGGGGATCCGCGCGCGGATCTAGTAAC | TGAAGACCATGGCGCGCCATGGTGGAGC |
| | pL4 | | TCTGGCGTAATAATTCGGGGGATCTGGATTTTAG | CAGGTCGACTCTAGAGGATCGTTTAAACGATCTGTCGATCGACAAGCTCGA |
| | pL5 | | GAGCTCGGTACCCGGGGATCGGCGCGCCATGGTCTTCACACTCGAAGATTTCG | CCCCGAATTATTACGCCAGAATGCGTTCGC |
| | pL6 | | tatagatgacaccgcgcgCGCGCGCGgatctagtaaca | gctcaagctaagctaagcttGTTTAAACgatctgtcgatcg |
| | pL7 | | CTCCACCATGGACCCACACATCTTTAATCGCC | GAAGACCATGGCTGGGCGGGCGGGGACCG |
| | *EcNIT4A2* promoter | | GCTGAGTTTGGTGGTGAAAA | CGACCTCGGCGATCAA |
| PCR amplification for Sanger sequencing-based *E* | EcNIT4A2-NOS-T | | CTCACCCTGTTGTTTGGTG | CCTTATATTTTCCAGCCATAGAAG |
| PCR amplification for Sanger sequencing-based genome | *OsNIT4A* | | AGCTTCAGCTCCTCCCCCACCAT | ACTAGCTGCAGCACATTGGTGATTTCTGACT |
| | *OsNIT4B* | | AGTACGACAAGTAGCACACGGCCT | TCGCGGGTGATACTAGCTGCAGCA |
| PCR amplification for Sanger sequencing-based hybridization | *EcALS3* | | CCCCAATCCCCCATCCAT | GCACCGCTCGCTGAATAC |
| Fine mapping | H001 | | GGTGGTGCTACGGAGACATAAT | GCGTTTCCAGTGTGATCACATC |
| | H005 | | TTCTTCTTGCTTTTCCGCTTCG | TCCTATGCCACATCAGCACATT |
| | H008 | | GCCAGATTGTACGTCGAAATGG | TGTGTTGCCTTCCTCTGTTCTT |
| | H009 | | ATGGAGGACAATCAACATGGCT | CTCAGTTCTCCCCTGGCTTTAG |
| | H011 | | ATACATGTGGGTAGGGTGGTGT | GACGTGACGATCGAGTAGCTAG |
| | H031 | | GCCGGAATCATAACTGCATGTC | TGAATGCCCTTTGTTATCCGGT |
| | H043 | | CCTATCAGGTGACATTGGGGAG | TTGACAAGCTGGCAATTCGTTT |
| | H045 | | CTGATTCCTCCCGTCGTCTCTC | GTGTCGACGCGGCTTAATTG |
| | H046 | | TGAAAAGGCAGAACACAACACC | CTGGTCCGTCCGATCTTTACAT |

Supplemental Table 2 *NIT4* genes used for sequence logo analysis

| Species | Gene ID |
|---|---|
| *Andropogon gerardi* | AndgeH1.04AG193900 |
| *Andropogon gerardi* | AndgeH1.04CG198500 |
| *Andropogon gerardi* | AndgeH1.04BG213000 |
| *Brachypodium arbuscula* | Barbu.4G136600 |
| *Brachypodium hybridum* | Brahy.S04G0135100 |
| *Brachypodium hybridum* | Brahy.D03G0685500 |
| *Brachypodium mexicanum* | Brame.04PG275100 |
| *Brachypodium stacei* | Brast04G127800 |
| *Brachypodium sylvaticum* | Brasy4G309100 |
| Bradi3g49630 | Bradi3g49630 |
| *Chasmanthium laxum* | Chala.05G203200 |
| *Echinochlao crus-galli* | NIT4A1 |
| *Echinochlao crus-galli* | NIT4A2 |
| *Echinochlao crus-galli* | NIT4A3 |
| *Eleusine coracana* | ELECO.r07.2AG0138490 |
| *Eleusine coracana* | ELECO.r07.2BG0194150 |
| *Hordeum vulgare* | HORVU6Hr1G059730 |
| *Miscanthus sinensis* | Misin08G213500 |
| *Oryza sativa* | LOC_Os02g42350 |
| *Panicum hallii* | Pahal.A02668 |
| *Panicum miliaceum* | AJ81AG22580 |
| *Panicum miliaceum* | AJ81BG24890 |
| *Panicum virgatum* | Pavir.1NG031200 |
| *Panicum virgatum* | Pavir.1KG396800 |
| *Paspalum vaginatum* | Pavag04G205600 |
| *Pharus latifolius* | Phala.02G214600 |
| *Saccharum officinarum x spontaneum* | SoffiXsponR570.03Eg173300 |
| *Saccharum officinarum x spontaneum* | SoffiXsponR570.03Ag214400 |
| *Saccharum officinarum x spontaneum* | SoffiXsponR570.03Dg218500 |
| *Saccharum officinarum x spontaneum* | SoffiXsponR570.03Cg201900 |
| *Setaria italica* | Seita.1G245900 |
| *Setaria viridis* | Sevir.1G250200 |
| *Sorghum bicolor* | Sobic.004G225200 |
| *Stenotaphrum secundatum* | Ss_v1_chr01G02382 |
| *Thinopyrum intermedium* | Thint.17G0494100 |
| *Thinopyrum intermedium* | Thint.17G0234600 |
| *Thinopyrum intermedium* | Thint.18G0332100 |
| *Thinopyrum intermedium* | Thint.18G0332400 |
| *Triticum aestivum* | Traes_6BL_C9E94B924 |
| *Triticum aestivum* | Traes_6AL_60AFB444B |
| *Triticum aestivum* | Traes_6AL_16008D7B1 |
| *Triticum aestivum* | Traes_6DL_BBF78B02F |
| *Triticum aestivum* | Traes_6AL_721FA45E3 |
| *Triticum aestivum* | Traes_6BL_98D63690F |
| *Urochloa fusca* | Urofu.1G274400 |
| *Zea mays* | Zm00001d051214 |

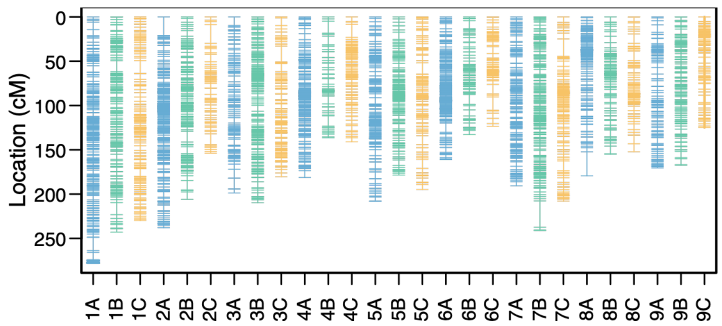


**Supplementary Figure 1 Linkage map of the S1 × R1 F2 population.**

Each vertical column represents one of the 27 chromosomes of hexaploid *Echinochloa crus-galli*, and horizontal ticks indicate the positions of individual markers. Genetic positions are shown in centimorgans (cM). The A, B and C homoeologous chromosomes are shown in blue, green and orange, respectively.

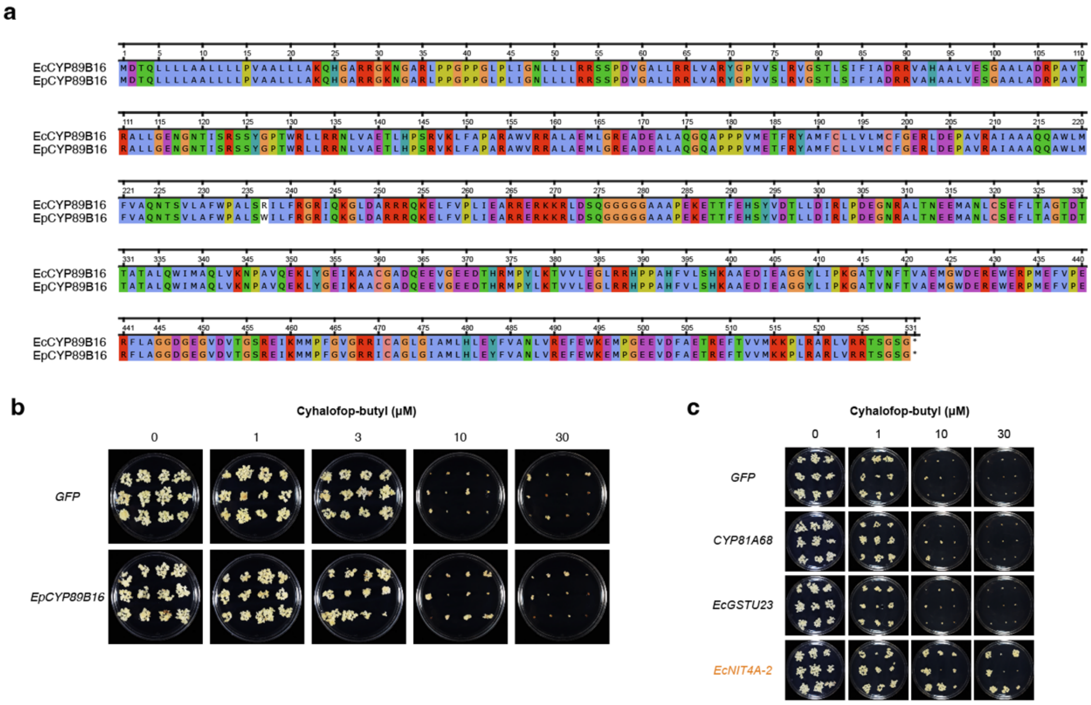


**Supplementary Figure 2 Cyhalofop-butyl sensivity of rice calli transformed with *EpCYP89B16*, *CYP81A68* or *EcGSTU23*.**

a) Alignment of EcCYP89B16 and EpCYP89B16. EcCYP89B16 and EpCYP89B16 differ only at amino acid residue 237. b,c) Calli 21 days after culture on cyhalofop-butyl-containing medium.

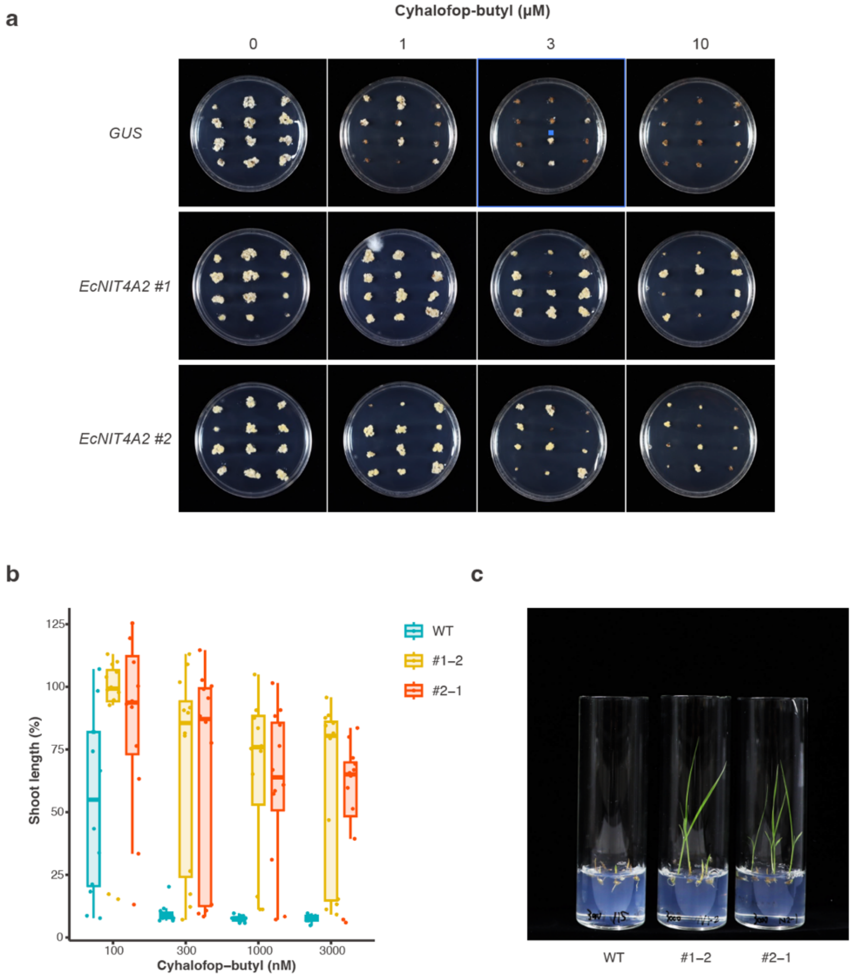


**Supplementary Fig. 3 Cyhalofop-butyl sensitivity of *EcNIT4A2*-expressing *Echinochloa oryzicola***

a, cyhalofop-butyl sensitivity of *E. oryzicola* calli expressing *EcNIT4A2*. Calli from two independent *EcNIT4A2*-expressing lines and two independent β-glucuronidase (GUS)-expressing control lines were cultured. Images were taken three weeks after transfer to the medium. b, shoot length of wild-type (WT) plants and two independent *EcNIT4A2*-expressing lines (#1–2 and #2–1) grown for 7 days on MS medium with cyhalofop-butyl. Each point represents a biologically independent seedling. Centre lines indicate medians, boxes indicate the interquartile range and whiskers extend to 1.5 times the interquartile range. c, Representative images of WT, #1–2 and #2–1 seedlings grown for 7 days on MS medium containing 3,000 nM cyhalofop-butyl.

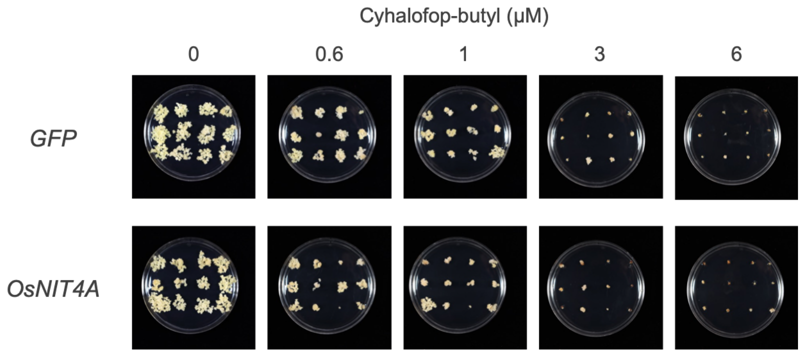


**Supplementary Figure 4 Cyhalofop-butyl sensivity of rice calli transformed with *OsNIT4A*.**

Images were taken three weeks after transfer to the medium.

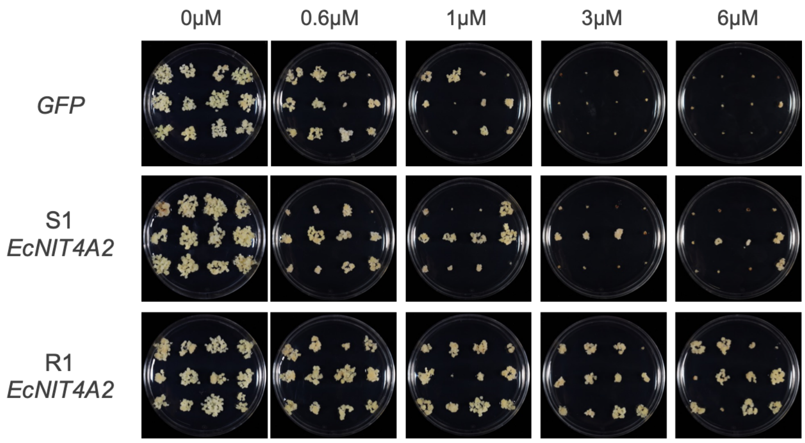


**Supplementary Figure 5 Cyhalofop-butyl sensivity of rice calli transformed with *EcNIT4A2* of S1 or R1 lines of *Echinochloa crus-galli*.**

Images were taken three weeks after transfer to the medium.

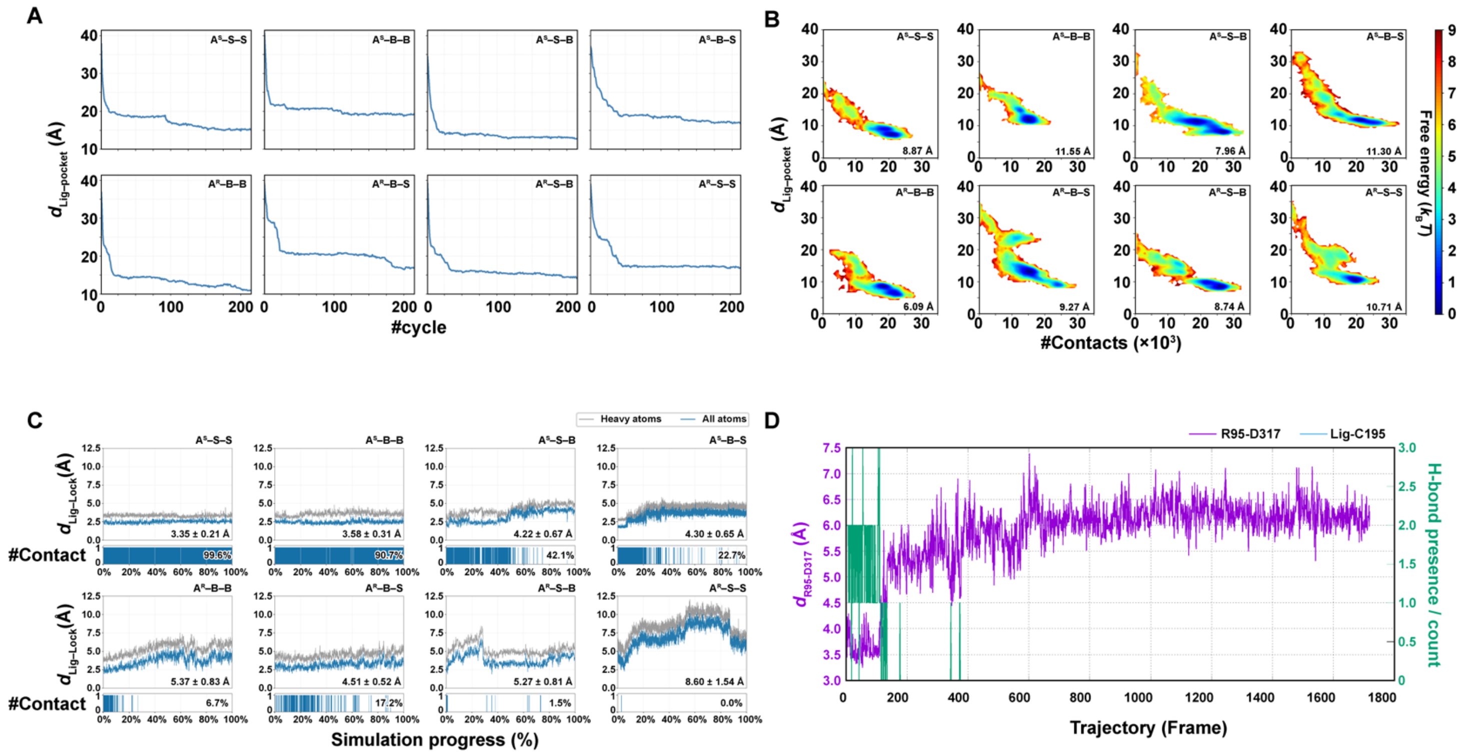


**Figure S6** LB-PaCS-MD analysis of cyhalofop-acid binding in eight EcNIT4A–EcNIT4B assemblies. A, Lid–pocket distances over 200 LB-PaCS-MD cycles for eight assemblies containing the $A^S$ or $A^R$ variant. B, Free-energy landscapes as functions of lid–pocket distance and the number of contacts, with minimum distances indicated in each panel. C, Ligand–lock distances calculated using all atoms (blue) or heavy atoms (grey) over normalized simulation progression. Lower strips indicate contact occurrence, with mean distances ± s.d. and contact percentages reported for each assembly. D, Time evolution of the R95–D317 distance (purple) and hydrogen-bond presence/count (green), with the ligand–C195 distance indicated in the legend. Assemblies are arranged in the same order in panels a–c, with $A^S$-containing systems in the upper row and $A^R$-containing systems in the lower row.

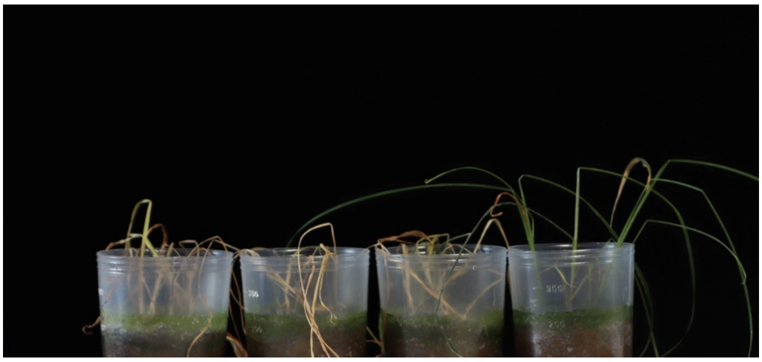

S4 S3 S2 R1

**Supplementary Figure S7 Cyhalofop-butyl sensitivity of *Echinochloa crus-galli* lines.**
Plant appearance 21 days after cyhalofop-butyl treatment.

Supplementary Video 1 Representative cyhalofop-acid binding trajectory for the D317 NIT4 system.

Supplementary Video 2 Representative cyhalofop-acid binding trajectory for the G317 NIT4 system.
